# Improved bacterial single-cell RNA-seq through automated MATQ-seq and Cas9-based removal of rRNA reads

**DOI:** 10.1101/2022.11.28.518171

**Authors:** Christina Homberger, Regan J Hayward, Lars Barquist, Jörg Vogel

## Abstract

Bulk RNA-sequencing technologies have provided invaluable insights into host and bacterial gene expression and associated regulatory networks. Nevertheless, the majority of these approaches report average expression across cell populations, hiding the true underlying expression patterns that are often heterogeneous in nature. Due to technical advances, single-cell transcriptomics in bacteria has recently become reality, allowing exploration of these heterogeneous populations, which are often the result of environmental changes and stressors. In this work, we have improved our previously published bacterial single-cell RNA-sequencing protocol that is based on MATQ-seq, achieving a higher throughput through the integration of automation. We also selected a more efficient reverse transcriptase, which led to reduced cell loss and higher workflow robustness. Moreover, we successfully implemented a Cas9-based ribosomal RNA depletion protocol into the MATQ-seq workflow. Applying our improved protocol on a large set of single *Salmonella* cells sampled over growth revealed improved gene coverage and a higher gene detection limit compared to our original protocol and allowed us to detect the expression of small regulatory RNAs, such as GcvB or CsrB at a single-cell level. In addition, we confirmed previously described phenotypic heterogeneity in *Salmonella* in regards to expression of pathogenicity-associated genes. Overall, the low percentage of cell loss and high gene detection limit makes the improved MATQ-seq protocol particularly well suited for studies with limited input material, such as analysis of small bacterial populations in host niches or intracellular bacteria.

**IMPORTANCE:** Gene expression heterogeneity among isogenic bacteria is linked to clinically-relevant scenarios, like biofilm formation and antibiotic tolerance. The recent development of bacterial single-cell RNA-sequencing (scRNA-seq) enables the study of cell-to-cell variability in bacterial populations and the mechanisms underlying these phenomena. Here, we report a scRNA-seq workflow based on MATQ-seq with increased robustness, reduced cell loss, improved transcript capture rate, and gene coverage. Use of a more efficient reverse transcriptase and the integration of a ribosomal RNA depletion step, which can be adapted to other bacterial single-cell workflows, was instrumental for these improvements. Applying the protocol to the foodborne-pathogen *Salmonella*, we confirmed transcriptional heterogeneity across and within different growth phases and demonstrated that our workflow captures small regulatory RNAs on the single-cell level. Due to low cell loss and high transcript capture rates, this protocol is uniquely suited for experimental settings in which the starting material is limited, such as infected tissues.

## INTRODUCTION

Until now, bacterial transcriptome studies have mainly relied on bulk RNA-sequencing (RNA-seq) (1). This approach provides averaged gene expression values across an entire cell population and therefore does not allow conclusions regarding transcriptional heterogeneity between individual bacteria. Yet, such phenotypic heterogeneity is a common microbial phenomenon (2). It is important for bacterial survival strategies such as bet-hedging, which allows fast adaptations to changing environments (3, 4) or biofilm formation, where individual cells take on highly specific roles within a community (5).

Dating to 2009, pioneering work established single-cell (sc)RNA-seq in eukaryotes (6). While this field rapidly evolved (7), the development of scRNA-seq in bacteria was slow to progress due to several challenges (8). Compared to eukaryotes, prokaryotic cells are much smaller, leading to less input material per cell. Single bacteria contain RNA in the femtogram range (9) and the average mRNA copy number is low, with only 0.4 copies / cell (10). Further challenges include efficient cell lysis, which is hampered by the bacterial cell wall, and capturing non-polyadenylated bacterial transcripts. These differences prevent a direct adaptation of most eukaryotic single-cell transcriptomic workflows.

Nevertheless, thanks to technical advances, bacterial scRNA-seq has recently become a reality (8). Three general types of approaches are currently available. Bacterial MATQ-seq (11), which stands for Multiple Annealing and dC-Tailing-based Quantitative scRNA-seq, is a workflow originally developed for eukaryotes (12) that relies on cell isolation by fluorescence activated cell sorting (FACS) and random priming of cellular transcripts. A second type, also previously established for eukaryotes and termed Split-Pool Ligation Transcriptomics Sequencing, or SPLiT-seq (13), is based on combinatorial barcoding. It was adapted for bacterial scRNA-seq in two independent studies introducing the so-called PETRI-seq and microSPLiT protocols (14, 15). In comparison to MATQ-seq, bacterial split-pool barcoding workflows enable the analysis of thousands of cells instead of a few hundreds, offsetting the lower transcript capture rate and higher rate of cell loss in these protocols. The third type is a microscopy- and probe-based approach that does not employ RNA-seq. It is called Parallel Sequential Fluorescence *In situ* Hybridization (par-seqFISH) and allows spatial transcriptomics on the level of single bacteria (16).

Despite these recent advances, challenges remain. These include a high frequency of cell loss and problems with robustness, coverage and prevalence of redundant ribosomal RNA (rRNA). In addition, short transcripts, such as regulatory small RNAs (sRNAs), show poor coverage or are not measurable at all. Importantly, transcript detection is currently limited to ~200 genes per single-cell, which is far below the average bacterial transcriptome. We reasoned that targeted improvements of our previous MATQ-seq protocol would address some of these challenges.

In this work, we present the next version of bacterial MATQ-seq. While the original protocol (11) has a high transcript capture rate, including low abundance transcripts, it is also limited in throughput and robustness. Through the integration of automation, we have now achieved increased cell throughput. In addition, we improved robustness through the selection of a more efficient reverse transcriptase (RT), which also led to a reduced transcript drop-out rate. Finally, given that the fraction of rRNA reads in our previous protocol reached up to 93%, we integrated a Cas9-mediated targeted rRNA depletion protocol, called Depletion of Abundant Sequences by Hybridization (DASH, (17)). This allowed us to obtain more gene expression information per single cell with decreased sequencing costs.

## RESULTS

### Automation of the MATQ-seq workflow achieves higher throughput

Initially, we aimed to increase cell throughput and read quality by integrating automation and by refining our analysis pipelines. In order to enable direct comparison with previous data, we performed all experiments in *Salmonella enterica* serovar Typhimurium. Within the MATQ-seq protocol, library preparation and quality control consist of a series of different labor-intensive pipetting steps. We implemented a user-friendly and highly flexible automation process by establishing protocols for all pipetting steps on the I.DOT dispensing robot (Dispendix), with the exception of clean-up and quality control steps (Fig. 1). This decreased turnaround times and the amount of consumables needed. Concurrently, automation increased sample throughput.

**Fig. 1.**
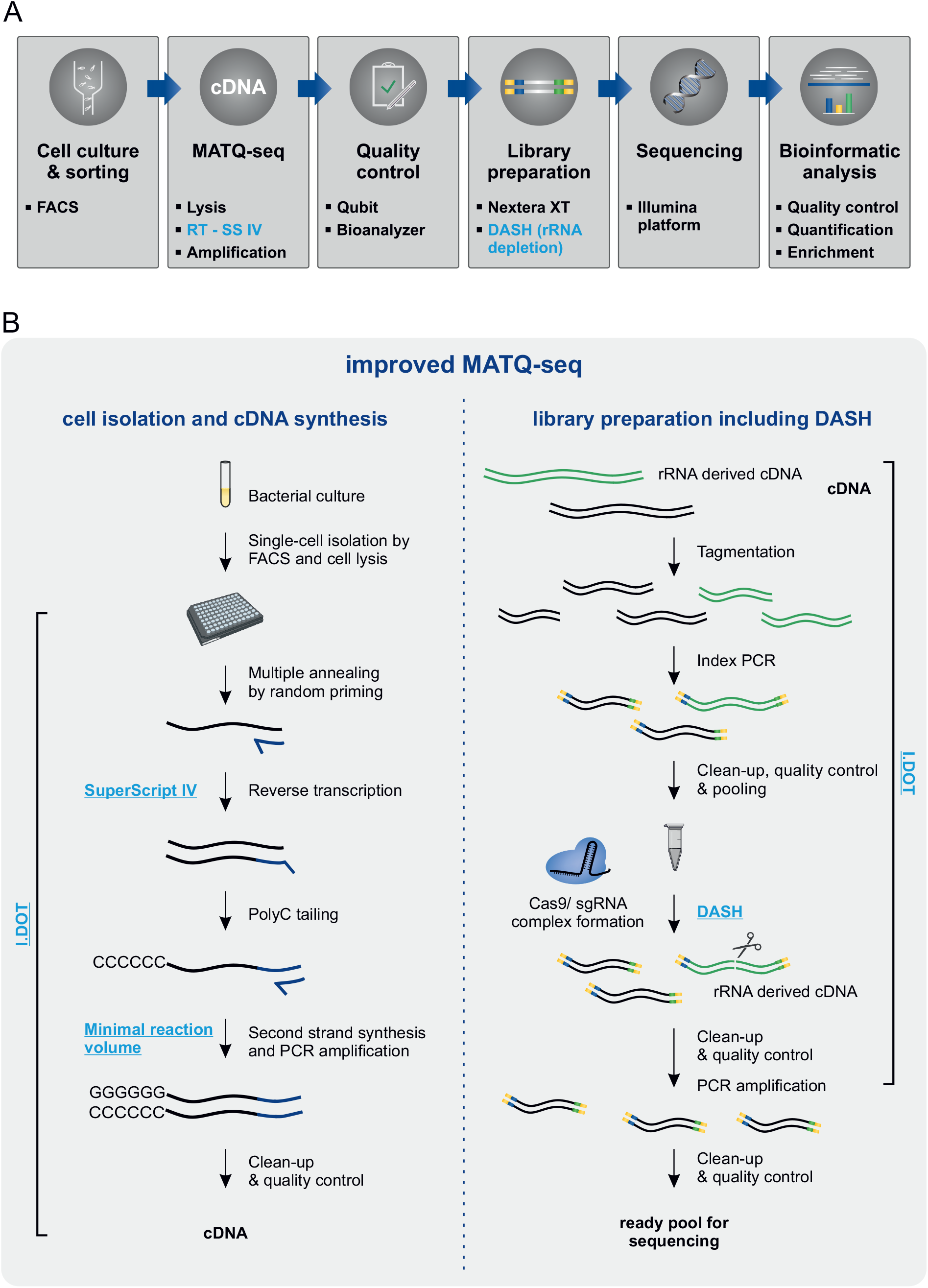
Improved MATQ-seq workflow for bacterial single-cell RNA-seq. **A)** Overview of bacterial scRNA-seq pipeline including major steps from cell culture to bioinformatic analysis. Changes compared to the previous MATQ-seq protocol are highlighted in blue. **B)** Detailed workflow of the MATQ-seq protocol separated into two main steps: (left) cell isolation and cDNA synthesis; (right) library preparation including DASH for ribosomal RNA depletion. Major improvements are highlighted in blue, including the use of SuperScript IV for reverse transcription, reaction optimization and integration of DASH into the library preparation. All pipetting steps were automated using the I.DOT dispensing robot, with the exception of all clean-up and quality control steps.

For cDNA analysis and quality control, we integrated the high-throughput Qubit Flex fluorometer for a precise and fast procedure. To facilitate sample processing, we applied a miniaturization step for the final PCR reaction volume by skipping the splitting step of the PCR reaction implemented in the original MATQ-seq protocol (12). This allowed processing of up to 96 single cells per MATQ-seq passage compared to a maximum of 24 cells in the previous protocol and decreased the overall processing time from about 10 to 8 hrs. More importantly, the hands-on time was reduced from about 6 to 3 hrs. Finally, we updated our data processing and analysis pipeline to improve data quality by implementing a better trimming approach, alignment, normalization, and identification of outliers (see Methods for details). Overall, all these steps discussed above led to higher cell throughput, improved accuracy and to higher read qualities, as described in more detail below.

### Optimized reverse transcription leads to higher robustness and reduced cell loss

Reverse transcription is crucial for RNA conversion and thus greatly affects the robustness of the scRNA-seq protocol and the detection of low-abundance transcripts. In our previous study, we used Superscript III (SS III) for the reverse transcription step (11). In the meantime, RTs with properties that promised to improve reverse transcription efficiency were reported (18–20). Therefore, we systematically tested Superscript IV (SS IV), an optimized RT based on SS III with improved thermostability, robustness and processivity; TGIRT and PrimeScript, two RTs that allow reverse transcription of GC-rich regions and highly structured RNAs; and Maxima H Minus and SMARTScribe, two highly efficient RTs with a high processivity and the capability to convert RNA transcripts up to 20 or 14.7 kb in length, respectively. For the analysis of these five different RTs we used fluorometer (Qubit) as well as high-resolution gel electrophoresis (Bioanalyzer) systems to assess cDNA yield and integrity.

Initially, we used total RNA as spike-in to evaluate the compatibility of the RTs with the MATQ-seq workflow (Fig. 2A). Of note, reverse transcription during MATQ-seq is performed with temperature gradients, which might interfere with RT efficiency. Indeed, SMARTScribe and PrimeScript showed low efficiency and were excluded from further validation. For the remaining three RTs we adapted buffer conditions to manufacturer’s recommendation. This yielded larger fragments in Bioanalyzer profiles, especially for SS IV and Maxima H Minus (Fig. S1A). Qubit measurements of cDNA obtained from spike-in tests using TGIRT showed low yield insufficient for further library preparation, ruling out this RT for further use (Fig. S1B).

**Fig. 2.**
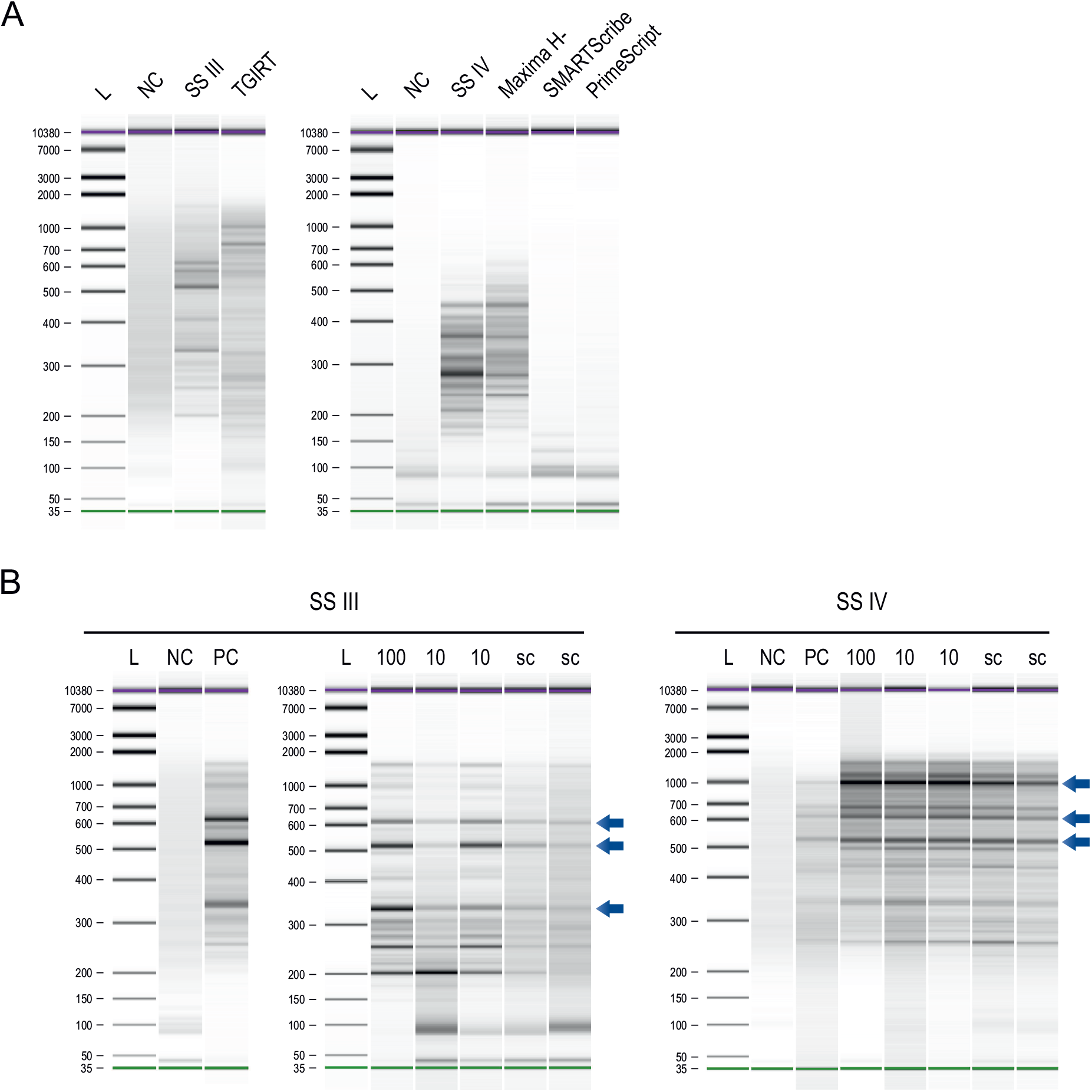
Selection of alternative reverse transcriptase. Bioanalyzer profiles of cDNA processed by MATQ-seq using different reverse transcriptases. **A)** Comparison of five different reverse transcriptases with Superscript III. Sample input was 50 ng of total RNA for all conditions. **B)** Comparison of cDNA profiles obtained with Superscript III (left panel) and Superscript IV (right panel). A gradient from 100, 10 and single sorted cells as input material is shown. The positive control was performed with a spike-in of 50 pg total RNA. Characteristic bands are indicated with blue arrows. L: ladder; NC: negative control; PC: positive control; sc: single-cell; SS III: SuperScript III; SS IV: SuperScript IV; Maxima H-: Maxima H Minus.

Next, we assessed the performance of SS IV and Maxima H Minus on sorted cells. Whereas both RTs showed similar results for 50 and 10 sorted cells, Maxima H Minus was less efficient on single-cell level (Fig. S1C). In a direct comparison of SS IV to SS III (Fig. 2B), SS IV showed a higher reproducibility, indicated by a pattern of characteristic bands and an increase in cDNA yield on the single-cell level. Based on these results and a lower cell loss overall, we implemented SS IV in the MATQ-seq protocol. Of note, due to the implementation of automation and the improvements in reverse transcription, only ~10% percent of all cells were lost during wet-lab processing (MATQ-seq and library preparation) and/or excluded by bioinformatic filtering.

### Ribosomal RNA depletion substantially increases non-rRNA reads

In order to reduce rRNA-derived reads, we applied an rRNA depletion step using DASH - a Cas9-mediated cleavage protocol originally developed in eukaryotes that can be implemented at the cDNA level (17). Specifically, a pool of single guide RNAs (sgRNAs) targeting rRNA-derived cDNA is provided together with Cas9, causing targeted cDNA cleavage. DASH had already been shown to work efficiently in low-input bacterial samples (>1 ng of total input RNA) (21) as well as on a single-cell level in eukaryotes (22).

In order to optimize DASH conditions to our protocol, we tested five molar ratios of Cas9:sgRNAs in the range from 1:2 to 1:50 and compared the percent of mapped rRNA reads (Fig. S2). A ratio of 1:2 was the most efficient and led to an rRNA depletion of 75%. These results are consistent with previously described DASH protocols for rRNA depletion in bacterial bulk samples, reporting depletion efficiencies in the range of 30-60% for *Salmonella* (21).

To evaluate rRNA depletion efficiency on a larger scale and across growth conditions, we applied it to *Salmonella* in early-exponential (EEP), mid-exponential (MEP), late-exponential (LEP) and early-stationary phases (ESP) (Fig. 3A). Per condition, 96 single-cells were processed by MATQ-seq. DASH is performed after tagmentation and introduction of full adapter index sequences by index PCR. Without this preamplification step, we were not able to recover enough cDNA after DASH. Since indices that allow cell identification are introduced during index PCR, samples can be pooled before DASH, thereby reducing the number of DASH reactions (Fig. 1, Fig. S3).

**Fig. 3.**
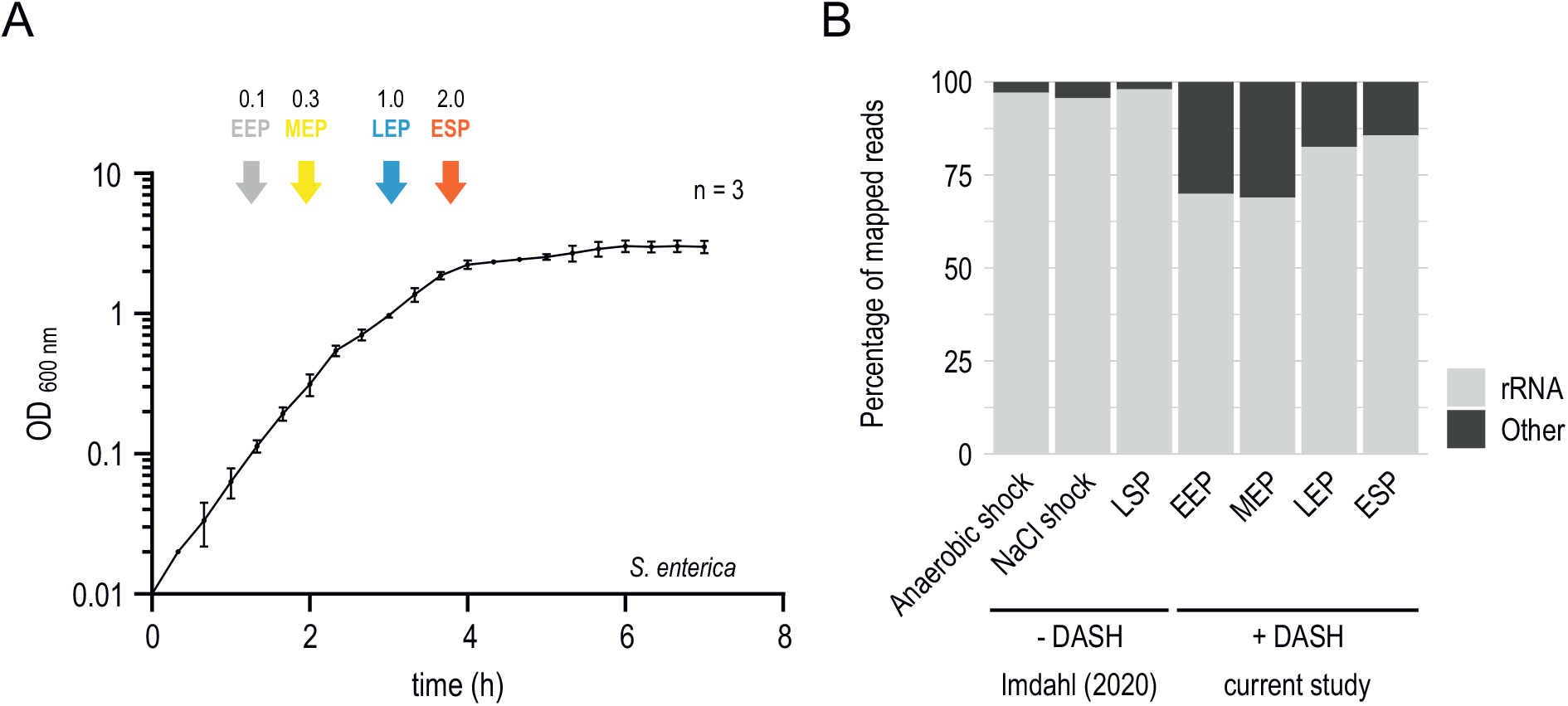
Experimental design and RNA class distribution. **A)** Growth curve of *Salmonella* in LB medium, with coloured arrows indicating the four sampling points for scRNA-seq experiments. Data are displayed as mean +/− standard deviation of three independent experiments. **B)** Representation of RNA class distribution comparing both protocols under different growth conditions. See Table S1 for detailed information on the prevalence of each RNA class. EEP: early-exponential phase: MEP: mid-exponential phase; LEP: late-exponential phase; ESP: early-stationary phase; LSP: late-stationary phase.

In comparison to data obtained using our original MATQ-seq protocol (11), the newly generated data set showed a much higher percentage of non-rRNA reads independent of growth condition (Fig. 3B). Importantly, in contrast to single-cell libraries that were not treated with DASH, we detected up to 10-fold higher percentage of reads mapped to coding sequences (CDSs), sRNAs, tRNAs and untranslated regions (UTRs), indicating successful elimination of rRNA-derived cDNA upon Cas9-mediated cleavage. Of note, the overall distribution of all other RNA classes did not vary substantially among growth conditions or between the two protocols, suggesting no major Cas9 off-target effect leading to unwanted cleavage of libraries (see Table S1). Compared to our initial test experiments (Fig. S2) using cDNA from sorted single-cells grown under anaerobic shock condition, the rRNA depletion efficiency was lower in this larger-scale experiment, although we used the same Cas9:sgRNA ratio. This might be due to an additional pooling step that could have saturated the Cas9 enzyme or to differences in commercial Cas9 batches that were used. Nevertheless, the DASH step still decreased rRNA reads up to ~30%.

The successful implementation of DASH required several adjustments to facilitate compatibility with the scRNA-seq workflow. Specifically, pre- and post-amplification cycles were optimized to ensure enough yield and, at the same time, to prevent over-amplification. We found that heat inactivation of proteinase K used in published DASH protocols was highly inefficient in our experimental settings. As a result, the downstream PCR was negatively affected by proteinase K and rRNA-depleted libraries were not amplified adequately. Inactivation by phenylmethylsulfonylfluorid (PMSF) led to higher efficiency and allowed PCR amplification of the final library pool (Fig. S3). Another important adjustment was the clean-up procedure required to remove PCR reagents, especially primer dimers from PCR products. The column-based purification used in published protocols was not suitable due to high sample loss. Instead, we used magnetic beads for the clean-up, which allowed us to purify low-input PCR samples with minimal sample loss. The ratio of magnetic beads and PCR product was adjusted to 1:1 (v/v), thereby ensuring capture of short fragments, including ones derived from short transcripts, such as sRNAs (Fig. S2).

### Improved MATQ-seq provides better gene coverage and shifts the gene detection limits

Due to the improvements we implemented, we were able to reduce the sequencing depth compared to our original MATQ-seq protocol (11) and still detect more genes on average across the four growth conditions compared to previous data: 307 versus 185 genes, respectively (Fig. 4A). We detected the highest number of genes in cells sampled in early-exponential phase (EEP) and mid-exponential phase (MEP), where more than 375 genes were detected on average (Fig. 4A). This is in line with expectations, as the mRNA level/cell has previously been observed to increase during exponential growth (23). Despite reducing the sequencing depth by ~6-fold, we achieved increased sensitivity of gene detection as the proportion of genes with no assigned reads (‘zeros’) across cells was reduced (Fig. 4B). This is directly related to the proportion of genes we detect across cells genome-wide, which increased to 95% (Fig. S4), but also genome-coverage, which increased from 3.0x to 4.8x (Table S2-3).

**Fig. 4.**
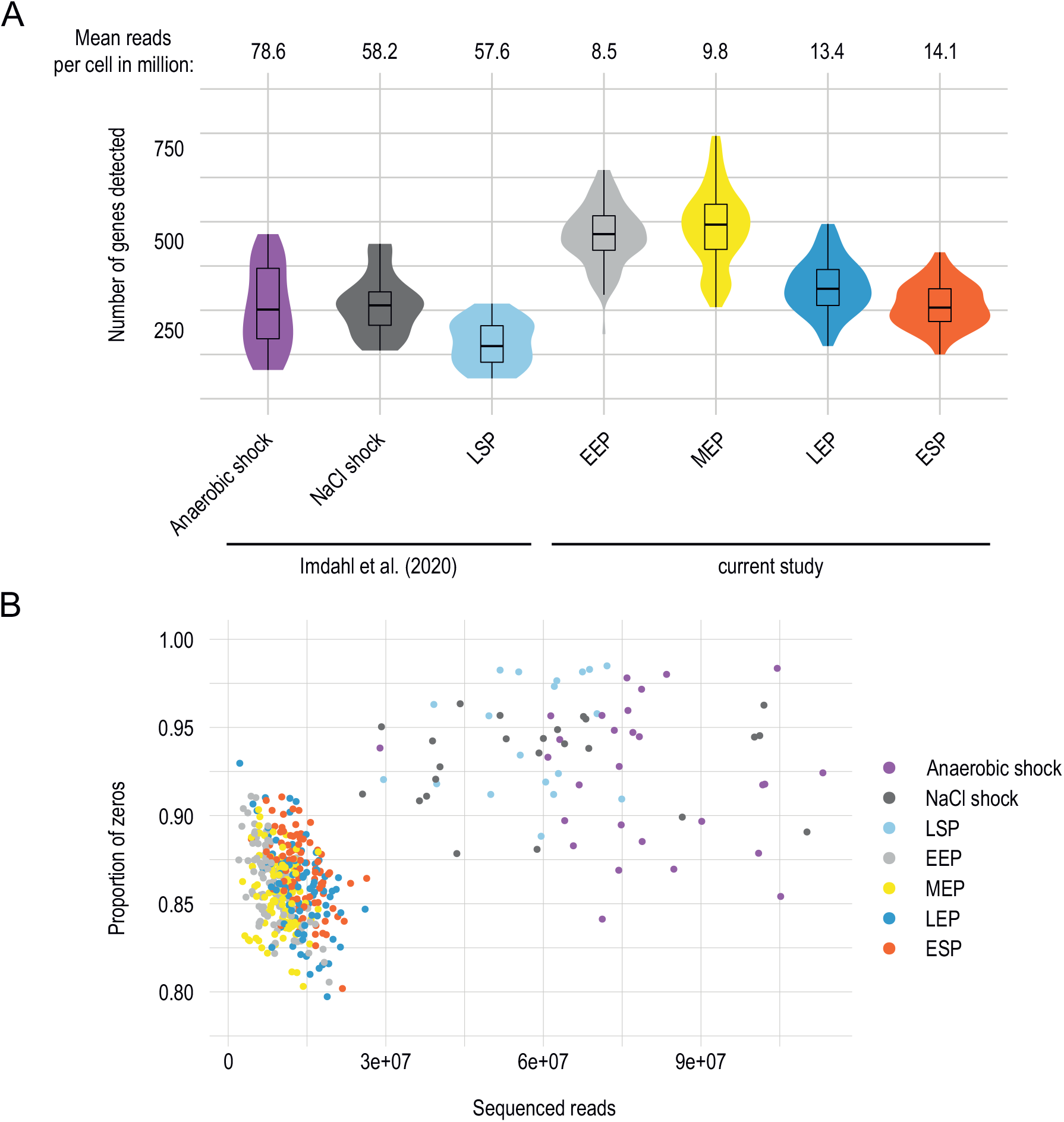
Gene detection limit and robustness of improved MATQ-seq workflow. **A)** Overlaid violin and boxplots showing the median, quartiles and distribution for the numbers of detected genes per condition. Mean numbers of reads per single-cell are indicated above. **B)** Proportion of genes with no assigned reads (‘zeros’) per single-cell compared to the number of sequenced reads, with each color-coded dot representing a single-cell.

We also wanted to assess how well our scRNA-seq data corresponded to condition-matching bulk RNA-seq data. Therefore, we generated pseudo-bulk data by summing gene expression across all cells per condition and compared it against bulk RNA-seq data of samples taken from the same culture as the sorted single-cells. Using three bulk RNA-seq replicates, we show higher correlations than previously observed (11), confirming a closer resemblance to bulk RNA-seq data (Fig. S5).

### MATQ-seq enables detection of small regulatory RNAs

sRNAs play a major role in bacterial gene regulation and are important during stress responses and virulence (24, 25). However, short transcripts, like sRNAs, are notoriously difficult to detect at the single-cell level due to inefficient recovery during the scRNA-seq workflow. In addition, clean-up procedures are required to remove primer dimers, but these bear risk to also target sRNAs which can be similar in size. Nevertheless, the use of magnetic beads at high ratio (1:1 v/v) that we adapted in MATQ-seq for this purpose improved the recovery of smaller fragments. As a result, we were able to detect a large number of sRNAs across the different growth conditions (Table S4), and between 28 to 41 unique sRNAs per condition (Fig. 5A). We observed a high variability in the overall prevalence and expression level of sRNAs across conditions, but also within the same growth condition (Fig. 5B). For example, the sRNAs CsrB and CsrC, known regulators of the global RNA-binding protein CsrA (26), were among the most prevalent sRNAs detected and showed highly variable expression across different growth conditions (Fig. 5B). The Csr complex constitutes one of the key regulatory systems for virulence, stress responses, motility and biofilm formation in *Salmonella* (27). In accordance with our data, condition-dependent differences in expression levels of both sRNAs have previously been described (28). The technical feasibility to detect sRNAs on a single-cell level was substantiated by *csrB* reads that only covered its transcribed region, indicating that coverage does not arise from processing artifacts. Though uneven mapping was observed among different single cells (Fig. 5C), a broad coverage of *csrB* transcripts suggests robust detection of sRNAs by MATQ-seq.

**Fig. 5.**
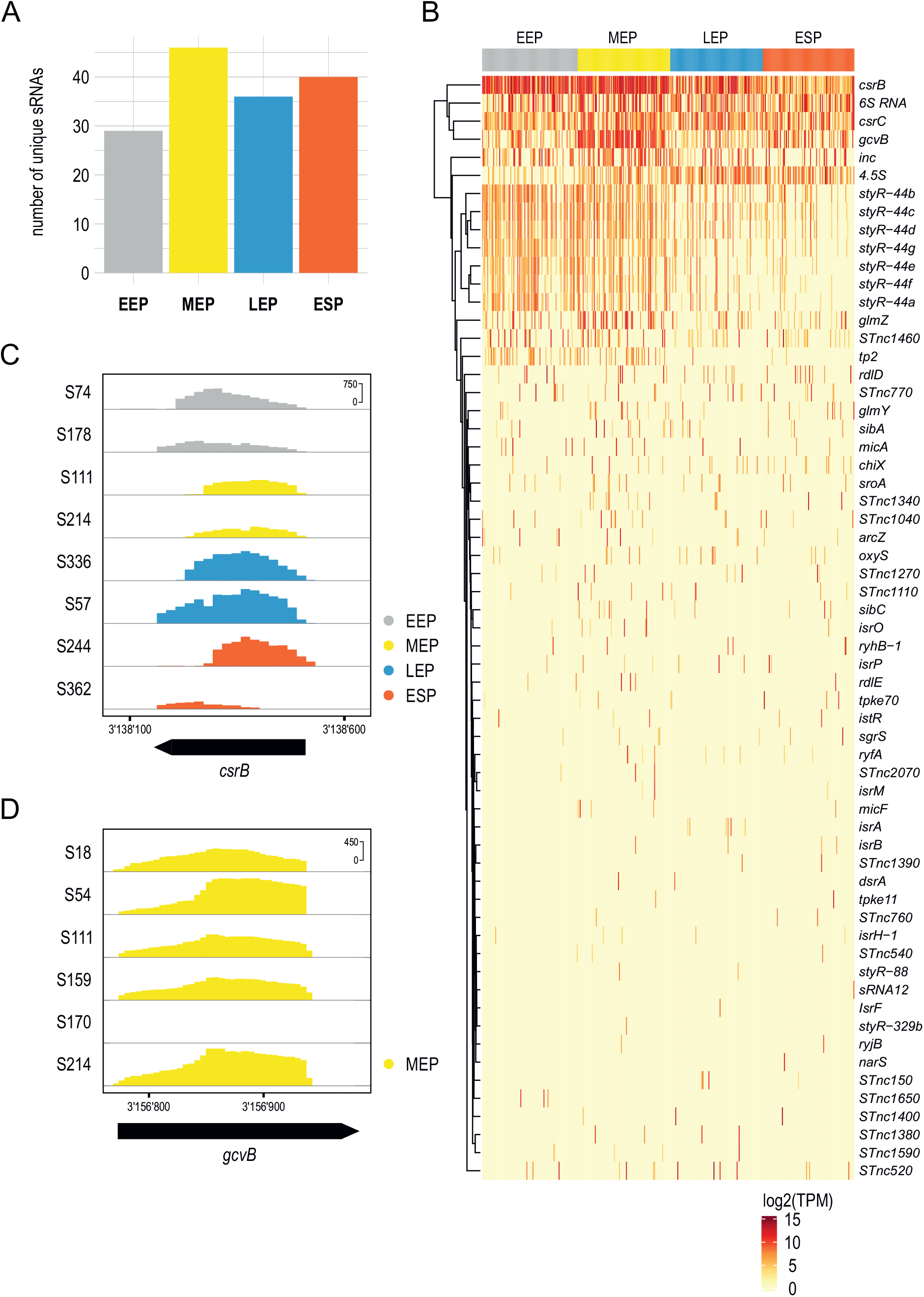
small regulatory RNA regulation on a single-cell level. **A)** Representation of unique sRNAs identified in each growth condition. **B)** Heatmap showing prevalence and distribution of the most abundant sRNAs in different growth conditions. **C)** Coverage plot and read densities of sRNA CsrB in eight selected single-cells (indicated by sample number). **D)** Coverage plot and read densities of sRNA GcvB in mid exponential phase (MEP) in six selected single-cells (indicated by sample number).

As an additional example, the sRNA GcvB, whose regulon mainly includes enzymes involved in amino acid biosynthesis and transporters, was highly expressed in the majority of cells in the MEP phase (Fig. 5B). Nevertheless we also observed cells that did not express *gcvB* (Fig. 5D). This finding is in accordance with earlier reports investigating *gcvB* expression in bulk samples; however, we did not observe a complete absence of GcvB in the stationary phase as previously seen (29). Instead, we detected a highly variable expression of GcvB in the ESP phase. Therefore, despite low (or non-detectable) expression on the whole population level, some cells still express GvcB, indicating heterogeneity across the cell population. Overall, these examples show that our improved MATQ-seq protocol enables detection of sRNAs on the single-cell level, reflecting expression patterns that have been previously reported using bulk RNA-seq data sets.

### Variability in Salmonella gene expression over different growth conditions

Visualization of all analyzed cells using principal component analysis (PCA) shows variation in gene expression over the different growth phases in *Salmonella*, as expected (Fig. 6A). EEP and MEP cells cluster together, in line with rapid cellular proliferation during these growth phases, which necessitates high gene expression (23). LEP and ESP are more distinct, reflecting the onset of nutrient starvation and a less active cellular state (30). Genes that drive the separation of these three main clusters are involved with growth-related processes and have previously been described (31). They include genes encoding components of flagella (*flaG*, also known as *flhB*, and *fliC*), lipid metabolism (*fadB*), glycolysis (*aceE*) and others (Fig. S6). Overlaying the gene expression across all cells in the same PCA plot helps visualize these expression patterns across and within conditions (Fig. 6B, C).

**Fig. 6.**
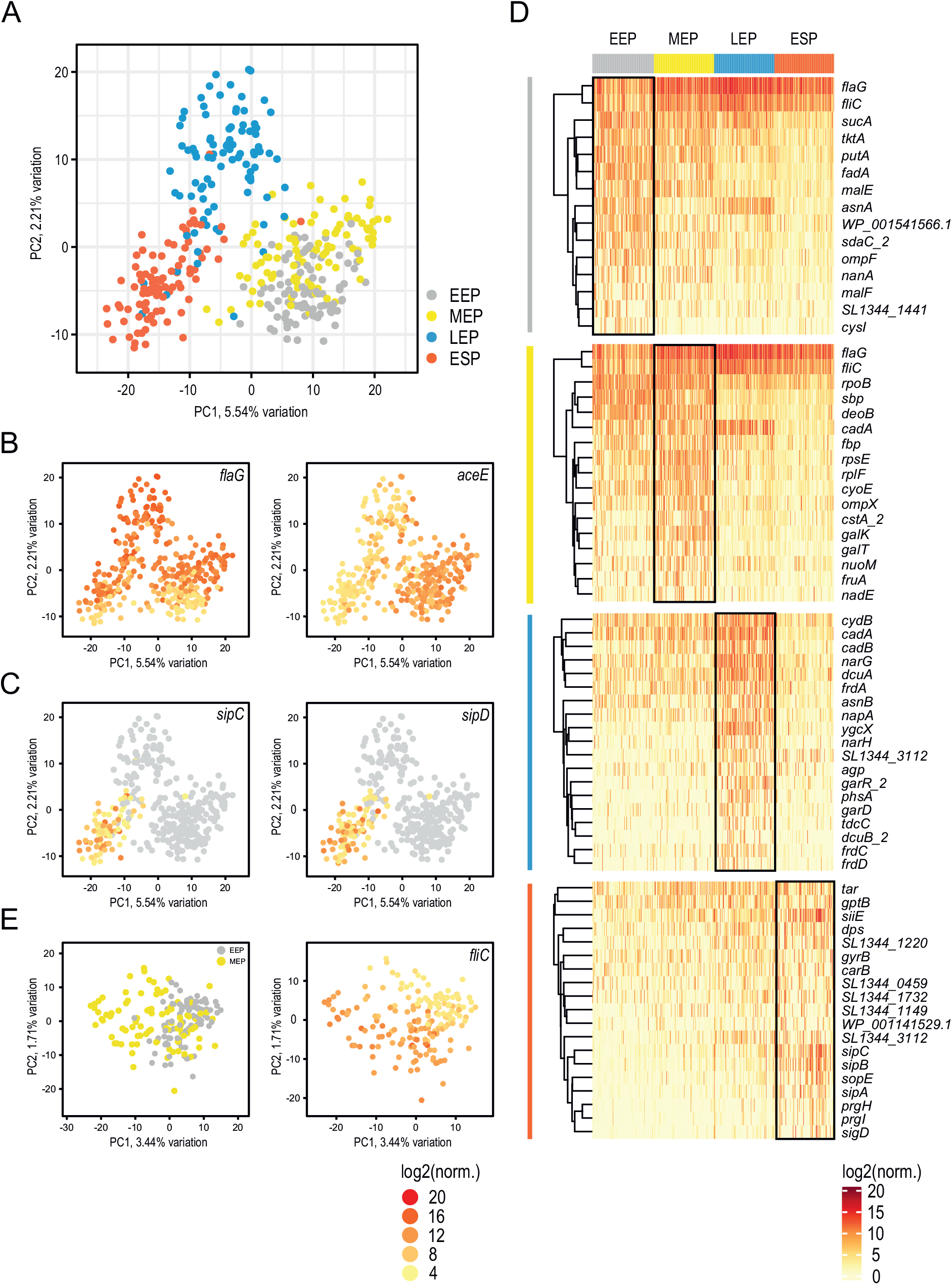
Cluster identification and analysis of highly variable genes detected on a single-cell level. **A)** Principal Component Analysis (PCA) of all analyzed cells across the four growth conditions. **B)** Overlay of the expression of genes contributing to the separation of the three main clusters seen in A. **C)** Expression of *Salmonella* pathogenicity genes *sipC* and *sipD* within early-stationary phase (ESP). **D)** Heatmap of the gene expression level of the top 10% of highly variable genes detected for each growth condition. **E)** (Left) PCA analysis of cells from early-exponential (EEP) and mid exponential phase (MEP). (Right) Overlay of expression of the flagella gene *fliC* on the PCA blot shown on the left.

We extracted the most highly variable genes (HVG) within each condition (Fig. 6D), highlighting their heterogeneous expression within a cell population. Of particular interest was the ESP phase, because we observed numerous genes related to *Salmonella* pathogenicity (*tar, siiE, sipA, sipB, sipC, prgH* and *prgI*) that appear to be associated with specific groups of cells (Fig. 6D). To explore this further, we focused on genes expressed from *Salmonella* pathogenicity islands (SPI) 1, 2 and 4 (Fig. S7). Here we observed three subpopulations based upon a group of ~7 SPI genes, exhibiting very low or no expression (middle), low expression (left) and high expression (right). Bulk RNA-seq studies have shown that SPI genes are more highly expressed during ESP compared to earlier growth conditions (31). Variation in expression of selected genes has previously also been reported at the single cell level (32). Here, by examining a larger set of SPI genes, we observe that a small subset of these genes mediates clustering of the cells into different populations.

To further analyze early growth conditions, we visualized only EEP and MEP cells with PCA (Fig. 6E). Due to the limited variation from each principal component, the cells remained closely clustered, further highlighting their similarities. Returning to the HVG, the two flagella genes *flaG* and *fliC* show particularly high variability in both growth phases (Fig. 6D). Both genes encode proteins of the core flagellum and their expression has a direct impact on cell motility (33). Overlaying the expression of *fliC* suggests two distinct subpopulations of cells, including both EEP and MEP cells (Fig. 6E, right panel). To visualize if additional flagella genes show the same pattern, we generated heatmaps containing flagella-expressing genes (Fig. S8). While we do not detect distinct subpopulations, we do see a gradient in flagellar gene expression, with a wide range of expression of flagella genes within each growth phase. This observation is in accordance with previous findings describing heterogeneous expression of *fliC* in *Salmonella* directly associated with cell motility and the potential to evade host inflammatory responses (2, 34).

### Effects of sequencing depth

The high number of sequenced reads per cell that we obtained allows us to explore gene detection limits per cell and how cell clustering is affected as sequence depth decreases. For both questions, we simulated scRNA-seq data across all cells for each condition separately (see Methods), querying different sequencing depths. Independent of the experimental condition, we reached saturation of the number of detected genes per cell at around 5 million reads (including rRNA-derived reads; Fig. S9A). This saturation analysis will assist future studies using MATQ-seq to find a balance between sequencing depth, the number of cells to analyze and the associated costs.

We also explored how PCA clustering is affected by the number of detected genes. For the four growth conditions tested, a minimum of 80 detected genes per cell led to qualitatively similar clustering results compared to using a much larger number of detected genes (Fig. S9B). Our expectation was that a greater number of detected genes would reveal more distinct sub-populations, but this does not appear to be the case within our experimental conditions, where a range between 80-126 genes appears to be sufficient to discriminate between growth phases. This suggests that it may be possible to investigate the structure of expression within a population with as few as tens of thousands of reads per cell given the efficiencies of cDNA conversion and rRNA depletion we have achieved here.

## DISCUSSION

In this study, we report substantial improvements to our previously established bacterial MATQ-seq protocol (11). Specifically, we focused on three elements of the workflow: (i) integration of automation and minimization of reaction volumes during different steps of the protocol, as well as optimization of the bioinformatic pipeline for data analysis; (ii) selection of a more efficient RT; (iii) implementation of an rRNA depletion step by integrating DASH into the library preparation. We validated our improved MATQ-seq protocol by generating a large data set of single *Salmonella* cells sampled over growth. Overall, our data show that the changes we implemented increased the cell throughput and robustness of the protocol, while reducing cell loss. In addition, we were able to improve gene coverage and the gene detection limits. We were even able to detect sRNAs on the single-cell level, which previously had not been feasible. This will allow the exploration of the regulatory functions of sRNA on the single-cell level in future studies. Moreover, our data confirm previously described heterogeneity within the same cell population, especially regarding *Salmonella* pathogenicity genes and genes encoding components of the flagellum (32, 34).

The successful implementation of DASH to deplete rRNA-derived cDNA was instrumental in achieving these advances. We believe that DASH can be adapted to other single-cell approaches, which currently do not include a targeted rRNA depletion protocol (11, 14, 15). Because depletion is performed at the cDNA level, DASH only needs to be customized to the library preparation step. For protocols that also use Nextera XT for library preparation, such as PETRI-seq (15), a direct application without any further adjustments is possible. The implementation of DASH would reduce the required sequencing depth and overall sequencing costs.

A general limitation of current scRNA-seq workflows is efficient cell lysis, which might require species-specific customization. Consequently, analysis of mixed bacterial communities is a challenge. This is especially true if their cell wall compositions vary, as this necessitates different enzymatic disruption and poses the risk of introducing bias based on varying lysis efficiency. Combinatorial indexing-based protocols (14, 15) can cope better with this limitation than MATQ-seq, because these protocols can process many more cells at once. Inefficient lysis is therefore compensated for by a higher number of input cells, although the danger of introducing bias remains. In contrast, MATQ-seq is more limited in throughput because of the cell sorting step and therefore inefficient lysis will lead to a high rate of failure. Nevertheless, MATQ-seq can in principle be applied to mixed cell populations if lysis conditions can be optimized.

Due to the high transcript capture rate of MATQ-seq, this method is particularly well suited for experimental settings in which the starting material is limited, such as the analysis of small subpopulations of bacterial cells in host niches or intracellular bacteria. In these settings, the application of Dual RNA-seq allows the study of host-pathogen interactions through the simultaneous analysis of the transcriptomes of both the bacteria and the host (1, 35). Single-cell Dual RNA-seq (scDual-Seq) has been attempted, but so far, bacterial gene detection has either been inefficient (36) or the experiments were performed under a high multiplicity of infection, which does not reflect physiological conditions (37). Since MATQ-seq was initially developed for scRNA-seq in eukaryotes (12), we see high potential in establishing scDual-Seq with MATQ-seq to capture both eukaryotic and prokaryotic transcripts. In this context it is interesting to note that DASH has been shown to remove bacterial as well as eukaryotic rRNA on single-cell level (22). Nevertheless, establishment of a scDual-Seq protocol based on MATQ-Seq will require further testing and validation.

## MATERIAL and METHODS

### Bacterial strains and growth conditions

The bacterial strains used in this study are listed in Table S5. *Salmonella enterica* serovar Typhimurium SL1344 (38) and constitutively GFP-expressing SL1344 strain (39) were grown in Lennox broth (LB) (tryptone 10 g/L, yeast extract 5 g/L, and sodium chloride 85.6 mM) medium. GFP-expressing *Salmonella* strain was only used for FACS gating. Bacterial cultures were inoculated to an optical density (OD) at 600 nm of 0.01 and incubated at 37°C with agitation (220 rpm) until early-exponential, mid-exponential, late-exponential and early-stationary phase (OD 0.1, 0.3, 1.0 and 2.0; according to (31)). 1 ml of each culture was pelleted and washed twice with 1 ml of 1 x Dulbecco’s phosphate buffered saline (1x DPBS). Afterwards the pellet was resuspended in 1 ml of 100 % RNAlater/ RNAprotect Tissue Reagent (Qiagen) and kept on ice. Right before sorting, samples are diluted 1:20 in 1x DPBS.

All pipetting steps in the sections below were automated using an I.DOT (Dispendix) dispensing robot except for clean-up and quality control steps.

### Isolation of single-cells

Isolation of single-cells was done as previously described by Imdahl et al. (11). Briefly, single-cells were isolated using a BD FACS Aria III for sorting individual cells into 96-well plates prefilled with lysis buffer (0.26 μl of 10×Lysis buffer (Takara), 0.03 μl of RNase Inhibitor (100 U/μl, Takara), 0.26 μl of DPBS (Gibco), 0.1 μl of Lysozyme (50 U/μl, Epicentre) and 1.95 μl of nuclease-free water (Ambion)). After sorting, plates were kept on ice and stored at −80°C until further processing.

### Improved MATQ-seq protocol

The improved MATQ-seq protocol is based on the protocol described by Imdahl et al. (11) with several modifications. Briefly, reverse transcription was performed using primers described by Sheng et al. (12). Instead of Superscript III, Superscript IV was used (Invitrogen) as reverse transcriptase without changing RT reaction volumes. As reaction buffer SS IV buffer was used instead. RT is followed by primer, RNA digestion and poly-C tailing. Subsequent Second Strand Synthesis was performed with only ¼ of reaction volume before PCR amplification. cDNA purification was performed using AMPure XP beads (Beckman Coulter) at a 1:1 v/v ratio. cDNA quality was checked by using Qubit Flex and 2100 Bioanalyzer DNA High Sensitivity kit (Agilent Technologies). Oligonucleotides used for MATQ-seq are listed in Table S6.

### DASH sgRNA pool generation

sgRNA pool for Cas9-based ribosomal depletion in *Salmonella* was generated according to Prezza et al. (21). Briefly, dsDNA template for *in vitro* transcription was generated using KAPA HiFi HotStart ReadyMix (Roche). After column-based Clean-up and quality control on Nanodrop and Bioanalyzer DNA 1000 Kit (Agilent), sgRNA pool was generated by *in vitro* transcription using MEGA Shortscript T7 Transcription Kit (Invitrogen). The final pool consisting of 797 sgRNAs was purified using the Monarch RNA Cleanup Kit (500ug, NEB). sgRNA quality control was performed on Qubit and Bioanalyzer RNA 6000 Pico Kit (Agilent).

### Single-cell RNA-seq: Library preparation including ribosomal depletion

cDNA obtained from single-cells after MATQseq was further processed for library preparation including ribosomal depletion protocol. Library preparation was done using the Nextera XT DNA Library Preparation Kit (Illumina) including DASH protocol (according to (21) with several modifications). Tagmentation was performed with only ¼ of reaction and 0.5 ng cDNA input according to manufacturer’s recommendations. Index PCR was performed with 13 cycles using IDT for Illumina Nextera DNA Unique Dual Indexes (Illumina). Obtained libraries were purified with AMPure XP beads (Beckman Coulter) at a 1:1 v/v ratio to ensure capturing sRNA-derived cDNA. After QC, up to 12 samples were pooled equimolar for ribosomal depletion. sgRNA/Cas9 complex formation is followed by DASH using the appropriate ratio of cDNA: Cas9: sgRNA. Cas9 enzyme was inactivated by Proteinase K (15 min at 37 °C). Afterwards, Proteinase K was inactivated by adding PMSF (1 mM final concentration). Depleted cDNA was purified by another round of AMPure XP bead clean-up and used as input for second PCR amplification. 0.5 ng depleted cDNA was used as input for Nextera XT reactions omitting the tagmentation steps. As primers i5 and i7 index-independent primers were used to amplify non-cleaved cDNA products. PCR was done with 13 cycles and clean-up was performed with AMPure XP beads at a 1:1 v/v ratio. Oligonucleotides used for second PCR are listed in Table S6.

### Total RNA extraction and library preparation

Bacterial RNA was isolated from *Salmonella* strain SL1344 grown under the same conditions as for scRNA-seq experiments. RNA extraction was performed with 1.8 ml of each *in vitro* culture using the TRIzol reagent (Invitrogen) according to the manufacturer’s recommendation. RNA quality was checked using Qubit RNA High Sensitivity Assay Kit (Invitrogen) and 2100 Bioanalyzer RNA 6000 Pico/ Nano kit (Agilent Technologies). Prior library preparation, DNAse treatment was performed using DNAse I kit (Thermo Fisher) followed by ribosomal RNA depletion. Ribosomal RNA (rRNA) was depleted using Lexogen’s RiboCop META rRNA Depletion Kit protocol according to manufacturer’s recommendation using 100ng total RNA as input per sample. DNA libraries suitable for sequencing were prepared using CORALL Total RNA-Seq Library Prep protocol (Lexogen) according to the manufacturer’s recommendation with 13 PCR cycles. Library quality was checked using a 2100 Bioanalyzer DNA High Sensitivity kit.

### Sequencing

Sequencing pools of single-cell as well as total RNA-seq libraries were checked using Qubit DNA High Sensitivity Assay Kit and a 2100 Bioanalyzer DNA High Sensitivity kit. Sequencing of library pools, spiked with 1% PhiX control library, was performed in single-end 100-cycle sequencing mode on the NextSeq 2000 or NovaSeq 6000 platform (Illumina). Demultiplexed FASTQ files were generated with bcl2fastq2 v2.20.0.422 (Illumina).

### Bioinformatics

#### Pre-processing

Read trimming and quality control of MATQ-seq reads was executed using BBDuk ((40)) and MultiQC (41). To efficiently remove primer and adapter sequences located at both ends of a read, we ran BBDuk in a two-pass procedure using the default adapter sequence database augmented with MATQ-seq specific sequences (Table S7). The first pass focused on the 5’ end, with parameters: *minlen=18 qtrim=r1 trimq=20 ktrim=1 k=17 mink=11 hdist=1 trimpolya=30;* while the second pass focused on the 3’ end with parameters: *minlen=18 qtrim=rl trimq=20 ktrim=r k=17mink=11 hdist=1*.

#### Read alignment and counting

Read alignment and counting was performed with Bowtie2 (42) and featureCounts (43), allowing a single mismatch and run-in *--local mode*. BigWig files were generated using deepTools (44), passing additional parameters: *--binSize 5 --normalizeUsing BPM*. We have employed the same gene detection method from the original MATQ-seq analyses (11), requiring a detected gene to have > 5 reads.

#### Normalization and differential expression

DESeq2 (45) was used for normalization and differential expression analysis, using size factors calculated by the *computeSumFactors* function in Scran (46) and other recommended parameters for DESeq2 single cell analysis, which included using the likelihood ratio test (LRT), *useT=TRUE, minmu=1e-6*, and *minReplicatesForReplace=Γnf*.

#### Identifying outlier cells

To identify outlier cells, we calculated the average number of detected genes per cell (> 5 reads) for each condition. Cells were determined to be outliers if their detected gene number varied more than two s.d above or below the mean, which removed 14 cells. One additional cell was removed based on the PCA plot generated using PCAtools (47).

#### Comparison with existing bulk and single cell RNA-seq data

To ensure fair comparisons between bulk RNA-seq and scRNA-seq datasets, we processed all fastq files using the same pre-processing, alignment and counting approaches as described above. For our pseudo bulk representation from the single cell data, for each condition, we summed the counts across all cells for each gene. Bulk RNA-seq were generated in parallel with the single-cell data.

#### Small RNA and highly variable genes

Additional small RNA (sRNA) were added to our annotation giving us 172 sRNAs in total. To show the most abundant sRNA in the heatmap for Figure 5, sRNAs were only shown if the rowsums of TPM normalized counts across all conditions was > 100. The full list of expressed sRNA are provided in (Table S4). Highly variable genes were identified using Scran (46), with the top 10% of HVG used for the heatmap in Figure 6. The *Salmonella* pathogenicity and flagellar genes used in the supplementary heatmaps (Fig. S7–8) were reduced to only show genes expressed in the examined conditions.

#### Single cell simulations and downsampling

Simulated data was generated using the *sample* function in R. All *Salmonella* genes (including rRNA) were sampled with replacement, with read count frequencies used as probability weights per cell for each condition. Different sample sizes were used to represent sequencing read depth and detected genes were the resulting uniquely sampled genes.

## Data availability

Gene Expression Omnibus (GEO) access upon request.

## Acknowledgments

We would like to thank Anke Sparman for helpful comments and editing the manuscript. We also acknowledge Esther Gottwald for great technical support on MATQ-seq workflow and library preparation; Laura Jenniches for discussions about simulating data; Emmanuel Saliba and Fabian Imdahl for help with MATQ-seq pipeline. We further thank the Core Unit SysMed at the University of Würzburg and the Sequencing Facility at HZI Brunswick for RNA-seq data generation.

This work was supported by a DFG Leibniz Prize to J.V. and the DFG-funded core unit MICROSEQ for microbial scRNA-seq (grant DFG INST 93/1105-1).

The authors have declared that no conflict of interest exists.

## Author contributions

C.H. performed the experiments, C.H. and J.V. designed experiments, R.J.H. and L.B. performed bioinformatic analysis, C.H., R.J.H., L.B. and J.V. wrote and edited the manuscript, L.B. and J.V. supervised research project.

## Supplementary Tables

Table S1: Proportion of mapped reads to gene biotypes

Table S2: Genome coverage new cells

Table S3: Genome coverage previous cells

Table S4: sRNAs - TPM expression

Table S5: List of *Salmonella* strains

Table S6: List of oligonucleotides

---

**Fig. S1.**
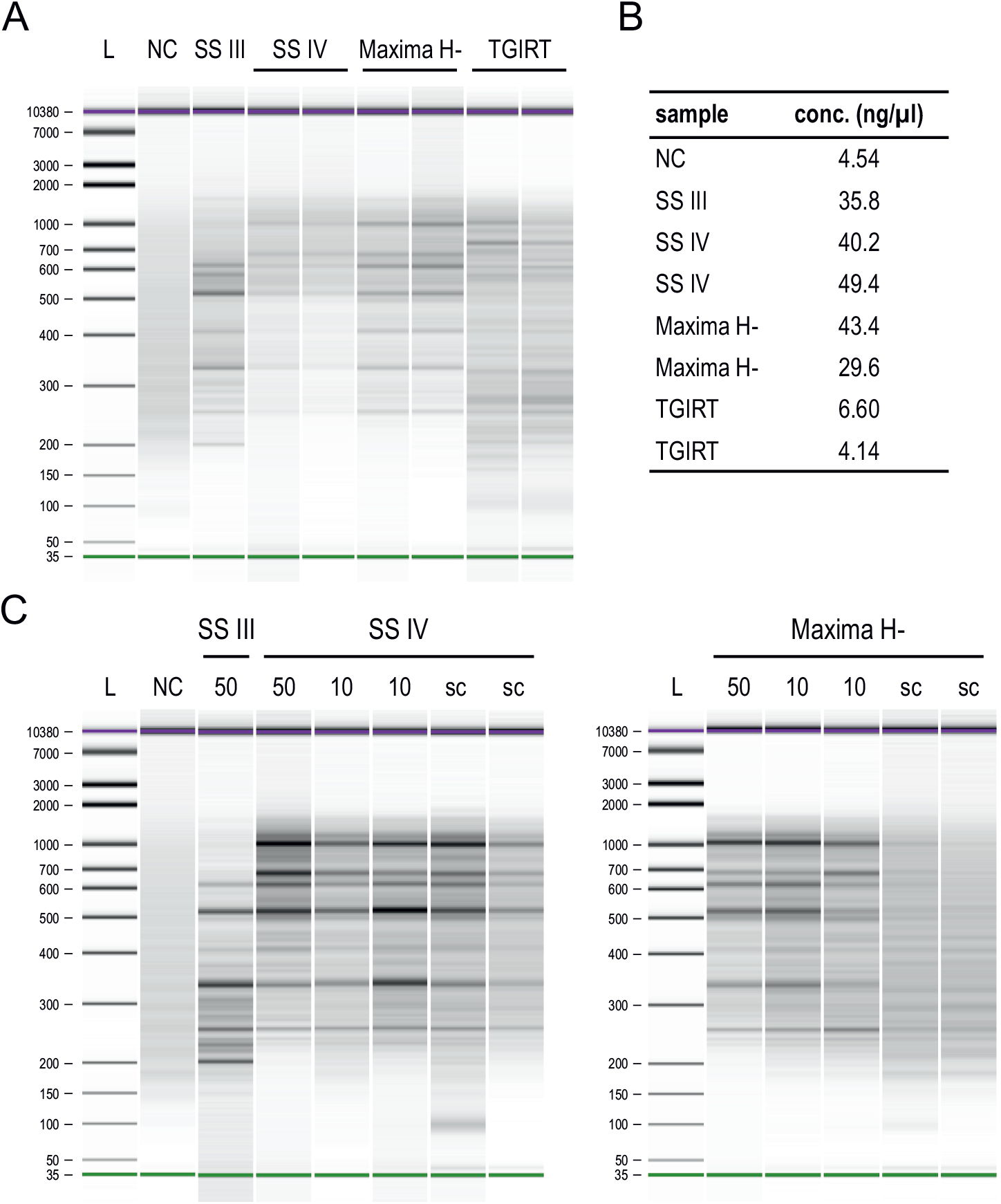
Validation of reverse transcriptases. Bioanalyzer profiles of cDNA processed with MATQ-seq using different reverse transcriptases. **A)** Further validation of the three RTs SS IV, Maxima H Minus and TGIRT after initial selection and subsequent buffer optimization (Fig. 2A). All assays were performed with a spike-in of 50 pg total RNA. **B)** cDNA concentrations of samples in A) measured with a Qubit fluorometer. **C)** Comparison of cDNA profiles obtained using SS III and SS IV (left panel) and Maxima H Minus (right panel). For SS IV and Maxima H Minus a gradient from 50, 10 and single sorted cells as input material is shown. L: ladder; NC: negative control; PC: positive control; sc: single-cell. SS III: SuperScript III; SS IV: SuperScript IV; Maxima H-: Maxima H Minus.

**Fig. S2.**
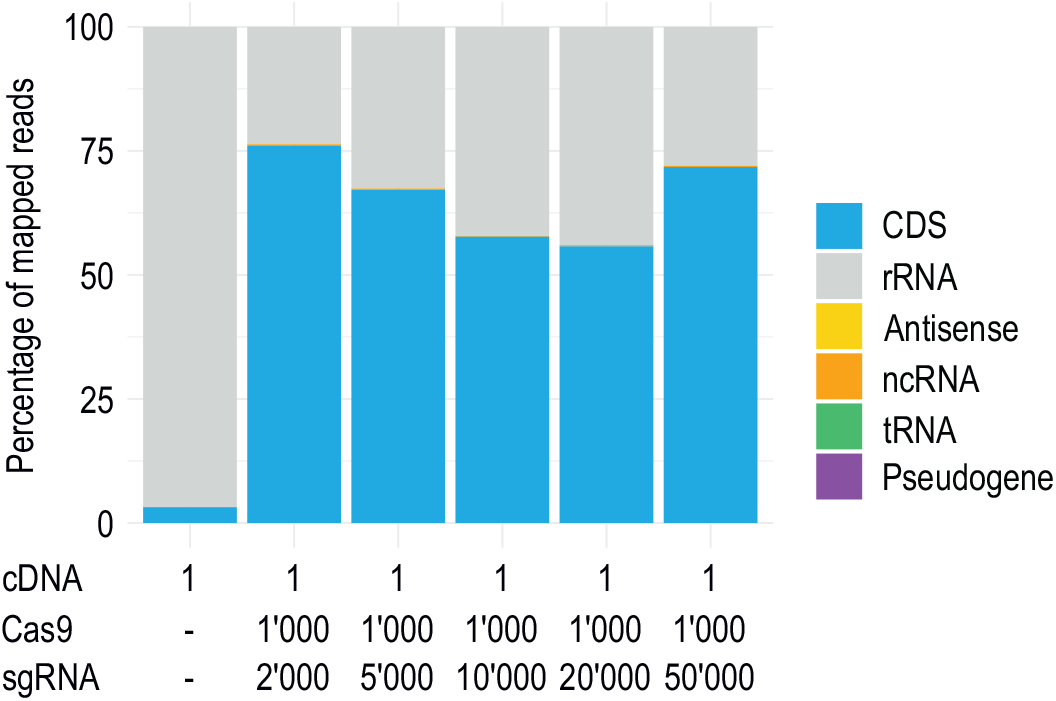
Comparison of different ratios of Cas9 and sgRNA for DASH. RNA class distribution obtained upon different Cas9:sgRNA ratios. Per ratio, three single-cells were processed by MATQ-seq including DASH under anaerobic shock condition. The RNA classes are shown as a percentage of the mean of the mapped reads (n = 3 single cells). Numbers below the graph represent the relative molar access of Cas9 and sgRNA over a single cDNA fragment. CDS: coding sequence; rRNA: ribosomal RNA; ncRNA: non-coding RNA; tRNA: transfer RNA.

**Fig. S3.**
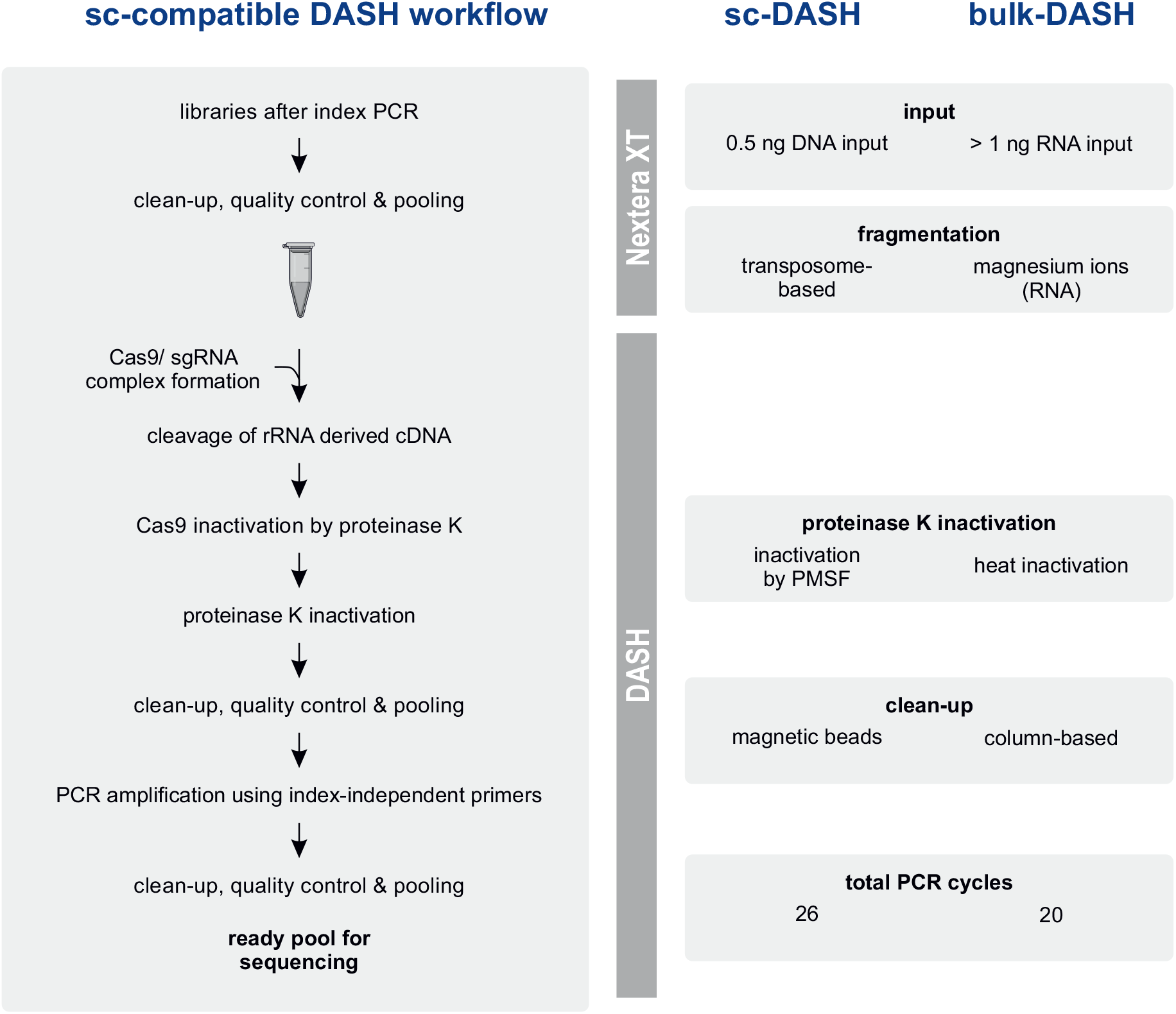
Implementation of sc-DASH workflow. Detailed overview of the integration of a single-cell compatible DASH workflow into the Nextera XT library preparation protocol. The right panel lists the most important adaptations required compared to the published bulk-DASH pipeline (Prezza et al. 2020).

**Fig. S4.**
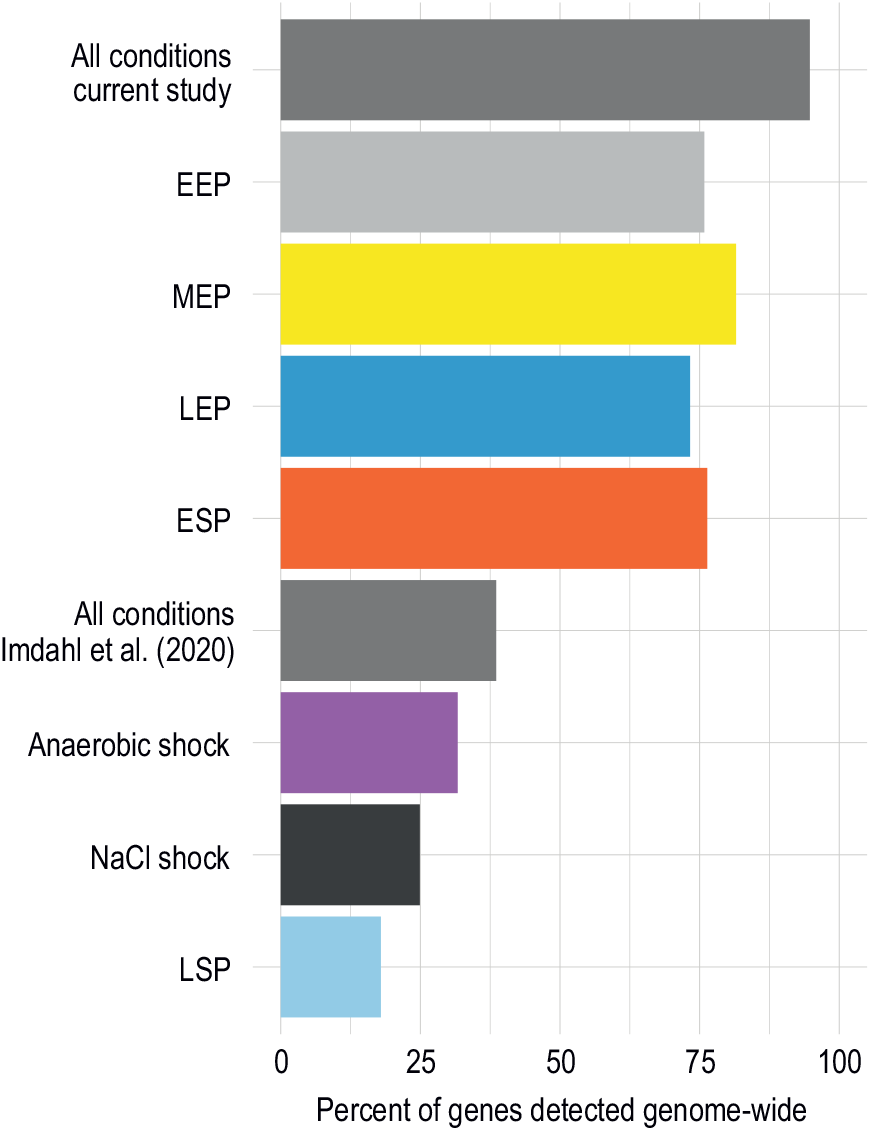
Percent of genes detected. The number of detected genes across all cells per condition for both MATQ-seq datasets.

**Fig. S5.**
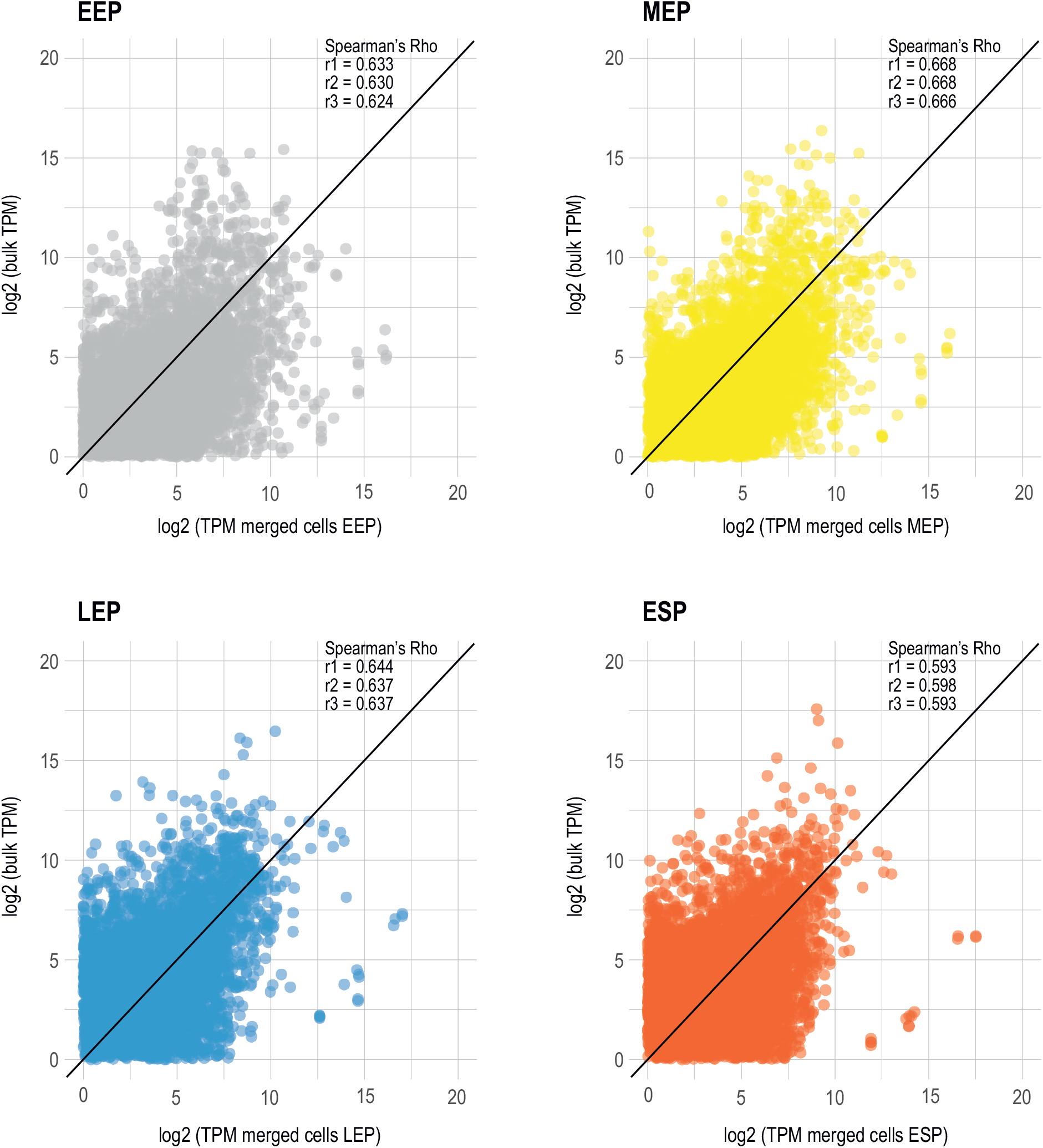
Correlations between scRNA-seq data and bulk RNA-seq data of different growth conditions. Spearman’s correlation between pseudo-bulk (counts summed across cells per condition) and bulk RNA-seq data for all four growth conditions; each dot represents a gene. Three bulk replicates were used and the associated correlations (r1 - r3) are shown. EEP: early-exponential phase: MEP: mid-exponential phase; LEP: late-exponential phase; ESP: early-stationary phase.

**Fig. S6.**
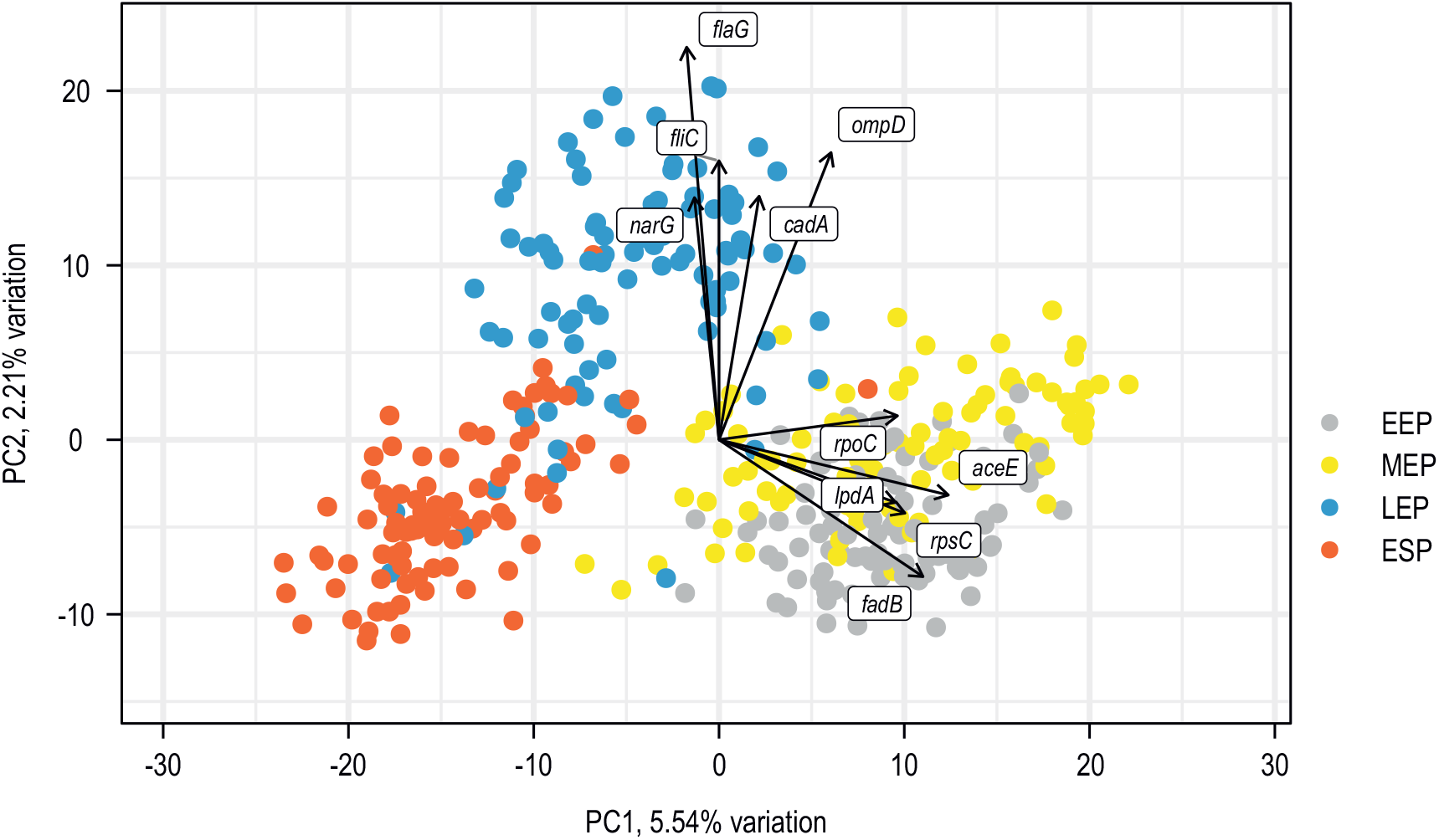
Drivers of heterogeneity. Principal component analysis based on Fig. 6A that includes the genes contributing the most amount of variation to separate the clusters across principal components 1 (x-axis) and 2 (y-axis). EEP: early-exponential phase: MEP: mid-exponential phase; LEP: late-exponential phase; ESP: early-stationary phase.

**Fig. S7.**
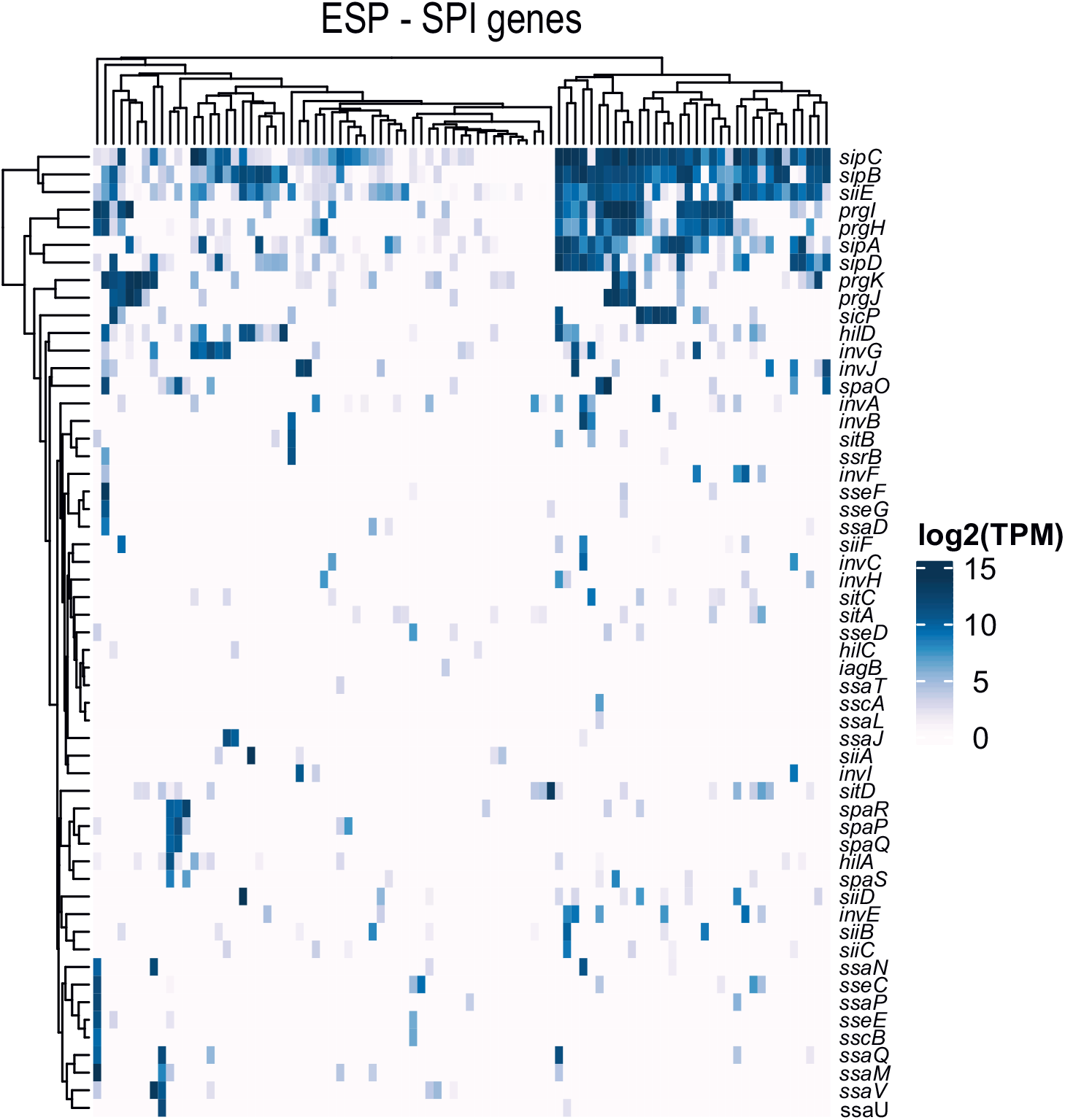
Heatmap SPI genes. Heatmap with cells from early-stationary phase (ESP) expressing *Salmonella* pathogenicity genes.

**Fig. S8.**
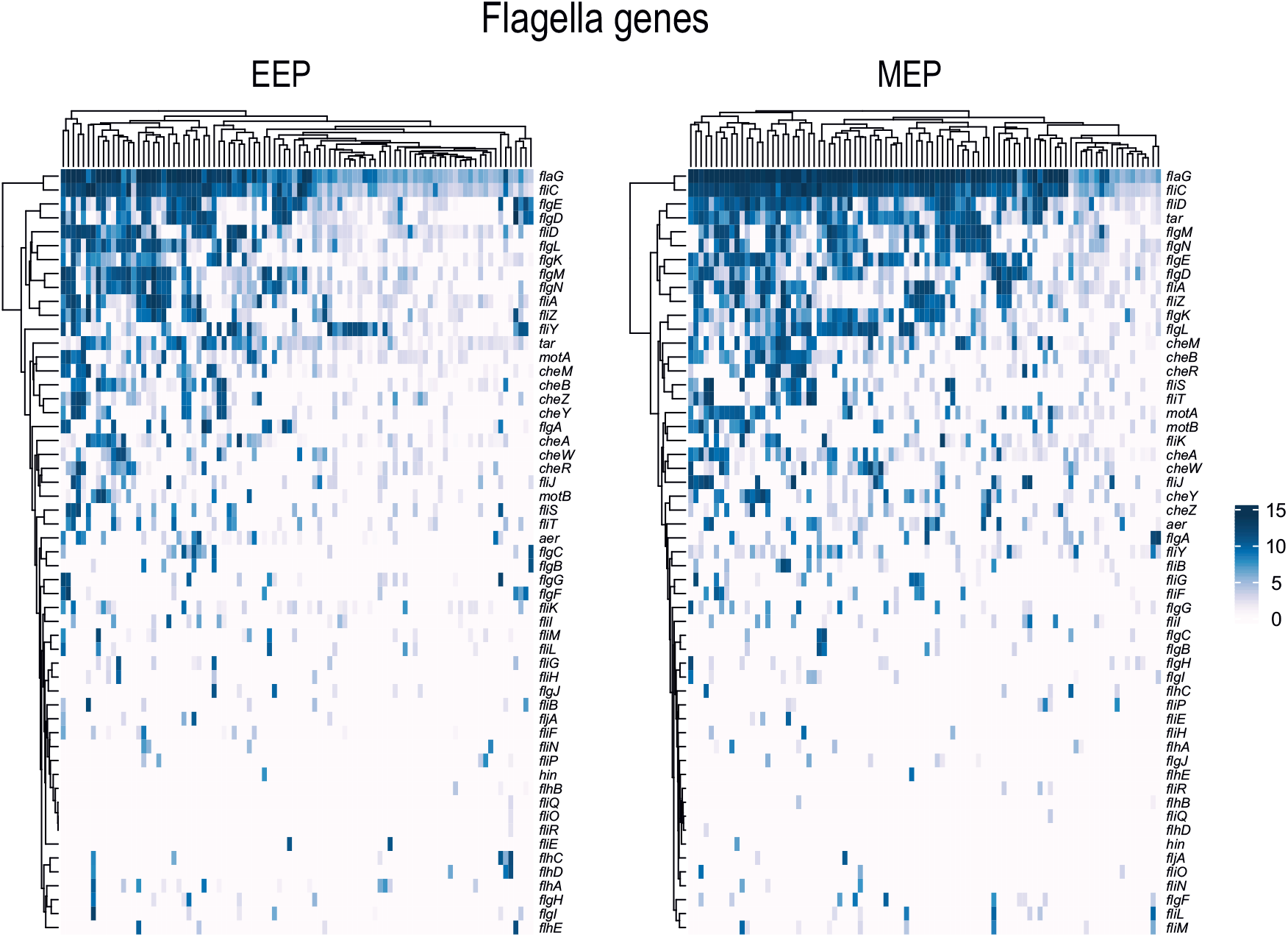
Flagella genes. Heatmaps with cells from early-exponential phase (EEP) and mid-exponential phase (MEP) expressing flagella genes.

**Fig. S9.**
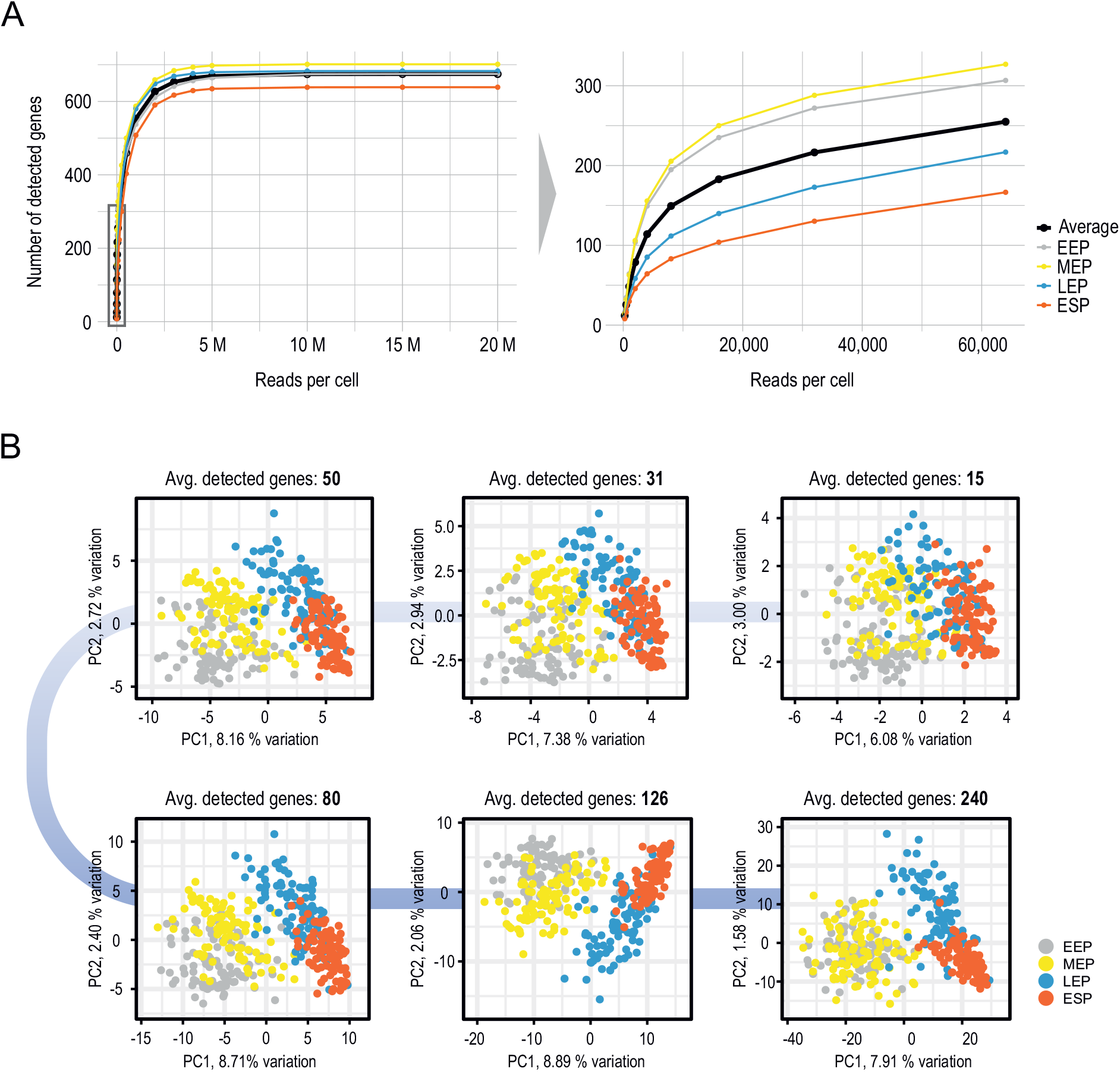
Saturation and downsampling PCA with detected genes. **A)** Reads per cell from simulated scRNA-seq data plotted against the number of detected genes. The right panel shows a zoomed-in view of the boxed area. **B)** Series of principal component analysis (PCA) plots from the same simulated data based upon different numbers of detected genes from 15 (top right) to 240 (bottom right).

